# The population genomics of archaeological transition in west Iberia: Investigation of ancient substructure using imputation and haplotype-based methods

**DOI:** 10.1101/134254

**Authors:** Rui Martiniano, Lara M Cassidy, Ros Ó’Maoldúin, Russell McLaughlin, Nuno M Silva, Licinio Manco, Daniel Fidalgo, Tania Pereira, Maria J Coelho, Miguel Serra, Joachim Burger, Rui Parreira, Elena Moran, Antonio C Valera, Eduardo Porfirio, Rui Boaventura, Ana M Silva, Daniel G Bradley

## Abstract

We analyse new genomic data (0.05-2.95x) from 14 ancient individuals from Portugal distributed from the Middle Neolithic (4200-3500 BC) to the Middle Bronze Age (1740-1430 BC) and impute genomewide diploid genotypes in these together with published ancient Eurasians. While discontinuity is evident in the transition to agriculture across the region, sensitive haplotype-based analyses suggest a significant degree of local hunter-gatherer contribution to later Iberian Neolithic populations. A more subtle genetic influx is also apparent in the Bronze Age, detectable from analyses including haplotype sharing with both ancient and modern genomes, D-statistics and Y-chromosome lineages. However, the limited nature of this introgression contrasts with the major Steppe migration turnovers within third Millennium northern Europe and echoes the survival of non-Indo-European language in Iberia. Changes in genomic estimates of individual height across Europe are also associated with these major cultural transitions, and ancestral components continue to correlate with modern differences in stature.

**Author Summary:** Recent ancient DNA work has demonstrated the significant genetic impact of mass migrations from the Steppe into Central and Northern Europe during the transition from the Neolithic to the Bronze Age. In Iberia, archaeological change at the level of material culture and funerary rituals has been reported during this period, however, the genetic impact associated with this cultural transformation has not yet been estimated. In order to investigate this, we sequence Neolithic and Bronze Age samples from Portugal, which we compare to other ancient and present-day individuals. Genome-wide imputation of a large dataset of ancient samples enabled sensitive methods for detecting population structure and selection in ancient samples. We revealed subtle genetic differentiation between the Portuguese Neolithic and Bronze Age samples suggesting a markedly reduced influx in Iberia compared to other European regions. Furthermore, we predict individual height in ancients, suggesting that stature was reduced in the Neolithic and affected by subsequent admixtures. Lastly, we examine signatures of strong selection in important traits and the timing of their origins.

## Introduction

Ancient genomics, through direct sampling of the past, has allowed an unprecedented parsing of the threads of European ancestry. Most strikingly, longitudinal studies of genomewide variation have revealed that two major technological innovations in prehistory, agriculture and metallurgy, were associated with profound population change [1–5]. These findings firmly address the longstanding archaeological controversy over the respective roles of migration, acculturation and independent innovation at such horizons; migration clearly played a key role. However, this may not be universal and genomes from several important European regions and time periods remain unexamined. In particular, at the southwestern edge of Europe several aspects of the archaeology suggest that some querying of the emerging paradigm is necessary.

First, whereas dating and similarity of the Portuguese Neolithic sites to other Mediterranean regions point to a rapid spread of agriculture at around 5500 BC [6], local Mesolithic communities were sedentary, dense and innovative -producing, along with Brittany, the earliest Megalithic tradition [7]. These groups persisted for at least 500 years after the onset of the Neolithic [8].

Second, in the transition to metallurgy, the Tagus estuary region of Portugal was a source for innovation. The distinctive Maritime Beaker, a key component of the Bell Beaker Package, characterised by grave goods including copper daggers and archery equipment first emerged there during the first half of the 3rd millennium BC. The Beaker package subsequently spread through Western Europe, where it is thought to have met and hybridized with the Steppe-derived Corded Ware or Single-Grave culture [9,10]. It remains an open question whether the influx of Steppe ancestry into North and Central Europe [4,5,11] associated with Corded Ware, also had a third millennium impact in Iberia.

Third, modern Iberia has a unique diversity of language with the persistence of a language of pre-Indo European origin in the Basque region. Interestingly, the population of Euskera speakers shows one of the maximal frequencies (87.1%) for the Y-chromosome variant, R1b-M269 [12], which is carried at high frequency into Northern Europe by the Late Neolithic/Bronze Age steppe migrations [4,5,13], although its arrival time in Iberia remains unknown.

In order to investigate the nature of cultural progression at Europe’s south Atlantic edge we analyse genomes from 14 ancient Portuguese samples from the Middle Neolithic through to the Middle Bronze Age (4200-1430 BC). For broader context we also impute genomewide diploid genotypes in these and other ancient Eurasians and investigate ancient population structure and examine temporal change in individual height.

## Results

### Ancient DNA extraction, sequencing and authenticity

DNA was extracted from the dense portions of fourteen petrous bones [3] excavated from eight archaeological sites across Portugal (S1 Fig), dated from the Middle Neolithic (MN) and Late Neolithic/Copper Age (LNCA) to the Bronze Age (BA) (S1 Text). Genomic coverage obtained was between 0.05x-2.95x and endogenous DNA estimates ranged from 5.6% to 70.2% (Table 1). Data authenticity was attested to by post-mortem deamination at the end of reads (S2 Text, S2 Fig) and low contamination estimates; X-chromosomes in males gave an average of 1.3% (0-2.3%) (S1 Table, S2 Table) and mtDNA 1.07% (0-1.71%) (S4 Table).

**Table 1.** Summary of the samples sequenced in the present study.

| Sample | Archaeological Site | Archaeological Context | Final number of aligned reads | Endogenous % | Coverage | Sex | mtDNA haplogroup | Y-chromosome haplogroup |
| --- | --- | --- | --- | --- | --- | --- | --- | --- |
| LC41 | Lugar do Canto | Middle Neolithic | 56108047 | 51.04 | 1.05X | XX | U5b1 | - |
| LC42 |  |  | 143759395 | 70.18 | 2.95X | XX | H3 | - |
| LC44 |  |  | 100174945 | 63.83 | 1.98X | XX | H1e1a2 | - |
| LC45 |  |  | 4012535 | 15.66 | 0.07X | XY | - | I |
| CA117B | Cabeco da Arruda I | Late Neolithic/ Chalcolithic | 17719307 | 21.25 | 0.36X | XY | J1c1b | I2a1b |
| CA122A |  |  | 80203819 | 42 | 1.71X | XY | H1e1a | G2a2a1 |
| CM364 | Cova da Moura |  | 36493410 | 44.12 | 0.76X | XY | H | I2a1b |
| CM9B |  |  | 113150709 | 65.04 | 2.56X | XX | J1c1 | - |
| DA96B | Dolmen de Ansiao |  | 98314678 | 51.6 | 1.88X | XY | K1b1a | I2a1a1a1b |
| MC337A | Monte Canelas |  | 2789478 | 8.77 | 0.05X | XY | - | - |
| MG104 | Monte do Gato de Cima 3 | Middle Bronze Age | 67818796 | 13.3 | 1.19X | XY | U5b3 | R1b1a2a1a2 |
| TV32032 | Torre Velha 3 |  | 44506573 | 16.47 | 0.83X | XY | X2b+226 | R1b1a2 |
| TV3831 |  |  | 43125279 | 15.47 | 0.95X | XY | H1+152 | R1b1a2a1a2 |
| VO10207 | Monte do Vale do Ouro 2 |  | 13357369 | 5.59 | 0.24X | XX | U5b1 | - |

### FineStructure Analyses

In order to harness the power of haplotype-based methods to investigate substructure in our ancient samples, we imputed missing genotypes in 10 out of 14 ancient Portuguese together with 57 published ancient DNA genomes, choosing those with >0.75X coverage and using the 1000 Genomes phase 3 reference haplotypes [2,3,5,11,14–22].

Comparison of imputed variants from down sampled genomes with those called from full coverage has shown that this approach gives genotype accuracy of ∼99% in ancient Europeans and we confirmed this using four down-sampled genomes from different time horizons included within our analysis [3,19] (S5 Text, S5 Fig). We observed that lower minor allele frequencies (MAF) imputed less accurately and we subsequently filtered for a MAF < 0.05 (S5 Text, S6 Fig). This gave 1.5 million markers with phase information called across each of the 67 samples. With these we first used CHROMOPAINTER [23] to generate an ancestry matrix which was utilized by fineSTRUCTURE [23] to identify clusters (Fig 1). The 67 Eurasian samples divided into 19 populations on the basis of haplotype sharing which are highlighted in a principal component analysis (PCA) calculated from the coancestry matrix (Fig 1A). Geographical and temporal locations are shown for these also, where Fig 1B shows four populations of hunter-gatherers (HG) individuals, Fig 1C, three populations belonging to Neolithic farmers and Fig 1D other groups containing samples ranging from Copper Age to the Anglo-Saxon period.

**Fig 1.**
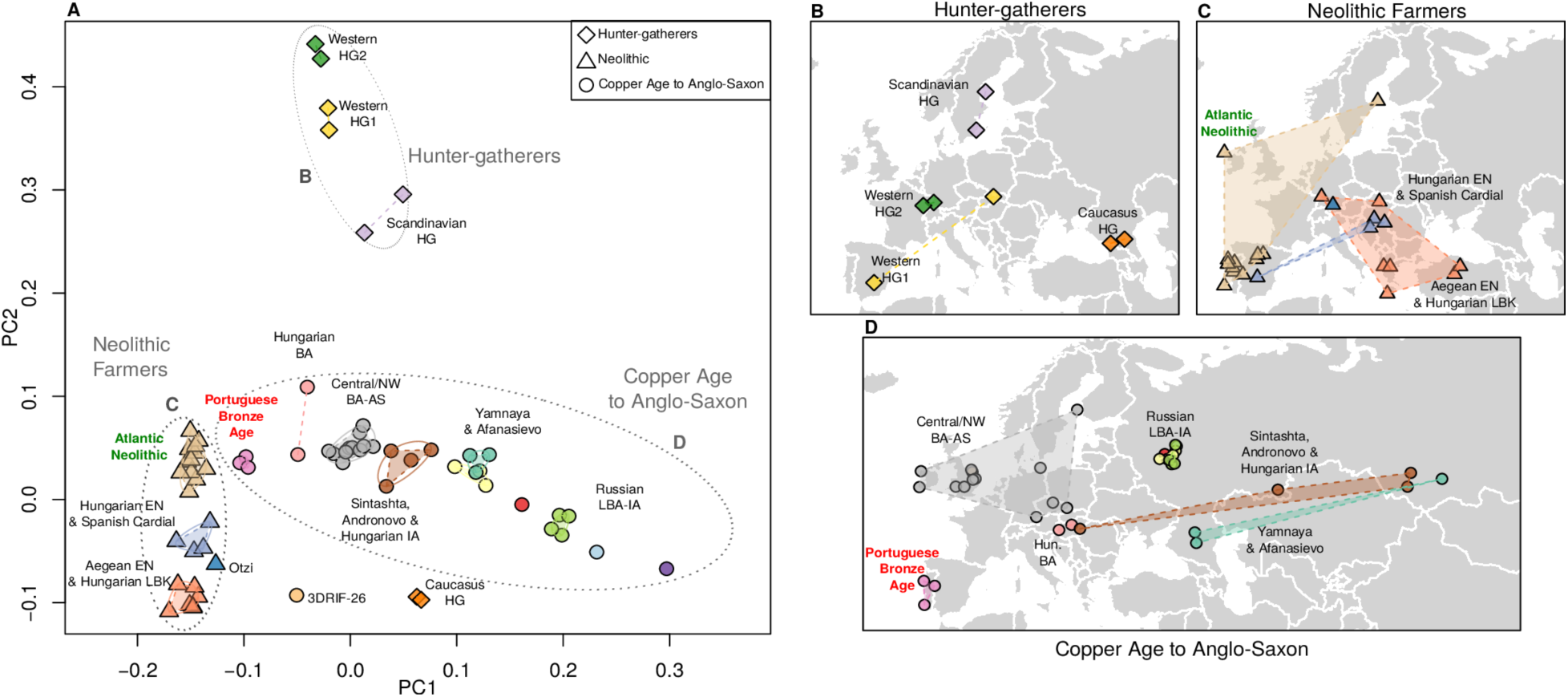
CHROMOPAINTER/fineSTRUCTURE Analysis. (**A)** PCA estimated from the CHROMOPAINTER coancestry matrix of 67 ancient samples ranging from the Paleolithic to the Anglo-Saxon period. The samples belonging to each one of the 19 populations identified with fineSTRUCTURE are connected by a dashed line. Samples are placed geographically in 3 panels (with random jitter for visual purposes): **(B)** Hunter-gatherers; (**C)** Neolithic Farmers (including Ötzi) and (**D)** Copper Age to Anglo-Saxon samples. The Portuguese Bronze Age samples (D, labelled in red) formed a distinct population (Portuguese*_BronzeAge*), while the Middle and Late Neolithic samples from Portugal clustered with Spanish, Irish and Scandinavian Neolithic farmers, which are termed “*Atlantic_Neolithic*” (C, in green).

Hunter-gatherer samples fall into 4 clusters (Fig 1B); interestingly the Paleolithic Bichon and Mesolithic Loschbour fall together (Western_HG1), despite 6,000 years separation, hinting at some level of continuity in the Rhine basin. Earlier Neolithic individuals are separated into two groupings, one comprising NW Anatolian and Greek samples, as well as two LBK individuals from Hungary and Germany. The second consists of Hungarian individuals from the Middle Neolithic to Copper Age alongside a Spanish Cardial Early Neolithic. A large cluster of individuals from Atlantic Europe, spanning the Middle Neolithic to Copper Age, is also seen, including all Portuguese MN and LNCA samples.

Samples belonging to the Copper Age and subsequent time periods in Russia showed strong stratification, with previous insights into ancient population structure in the steppe [5] corroborated by the formation of the Yamnaya_Afanasievo cluster and the Sintashta_Andronovo. In contrast, Central/Northern European samples stretching from the Copper Age to Anglo Saxon period all clustered together with no detectable substructure (*CopperAge_to_AngloSaxon*). However, the Portuguese Bronze Age individuals formed a distinct cluster. This was seen to branch at a higher level with the *Atlantic_Neolithic* rather than *CopperAge_to_AngloSaxon,* and in the PCA plot placed between the two.

### Increase in local hunter-gatherer ancestry in the Middle and Late Neolithic

It has been previously shown that an individual (CB13) dating from the very beginning of the Neolithic in Spain showed ancestry closer to a Hungarian hunter-gatherer (KO1, found within a very early European Neolithic context) than to the more western HGs from LaBrana in Spain and Loschbour in Luxembourg [18]. Furthermore, recent studies have highlighted an increase in western hunter-gatherer (WHG) admixture through the course of the Spanish Neolithic [17,24]. To investigate suspected local HG introgression in Iberia we compared relative haplotype donation between different hunter-gatherers within European farmers and other samples from later time-periods (Fig 2). In Iberia, a clear shift in relative HG ancestry between the Early Neolithic (EN) to MN was observed, with greater haplotype donation from the Hungarian HG within the Cardial Neolithic sample CB13 (Olalde et al., 2016), when compared to other HG of more western provenance (Bichon, Loschbour and LaBrana). A reversal of this trend is seen in the later Neolithic and Chalcolithic individuals from Portugal and Spain, but intriguingly not in other Atlantic Neolithic samples from Ireland and Sweden. This is confirmed by a Mann-Whitney test demonstrating that Iberian Neolithic samples receive significantly more (p=1.02x10^-6^) haplotypes from west European HG (Bichon, Loschbour and LaBrana) than KO1 relatively to Neolithic samples from elsewhere in Europe suggesting a more prolonged hunter-gatherer interaction at the littoral. In the transition to the Portuguese Bronze Age, a second shift can be seen in relative hunter-gatherer ancestry with some increase in relative haplotype donation from KO1, which is seen more prominently in the majority of post-Neolithic Eurasian samples, hinting at some difference between the Portuguese Neolithic and Bronze Age.

**Fig 2.**
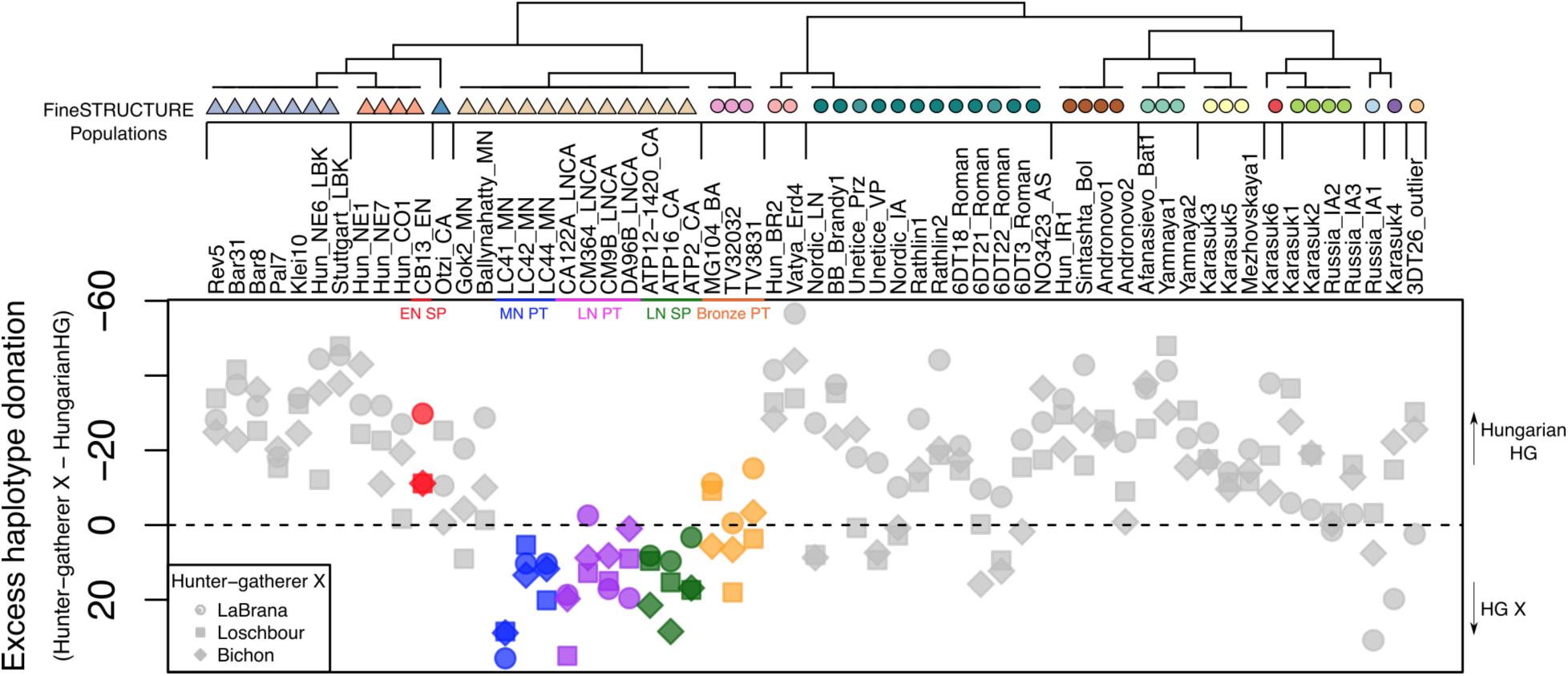
Patterns of hunter-gatherer haplotype donation to ancient Eurasians. This was estimated by subtracting the vector of haplotype donation of Hungarian HG from a vector of hunter-gatherer X, where X={LaBrana, Bichon, Loschbour}. Legend: E -Early; M -Middle; L -Late; N -Neolithic; PT -Portugal; SP -Spain. Note: HG individuals were removed from the tree.

### Steppe-related introgression into the Portuguese Bronze Age

Next, to further investigate this apparent shift between the Neolithic and Bronze Age in Iberia, we explored haplotype sharing patterns of ancient samples in the context of modern populations. We merged our dataset of imputed variants with 287,334 SNPs typed in 738 individuals of European, Middle Eastern, North African, Yoruba and Han Chinese ancestry [25] and ran CHROMOPAINTER/FineSTRUCTURE as above.

When comparing vectors of haplotype donation between Neolithic and Bronze Age individuals of different European regions to modern populations, a geographical pattern emerges (Fig 3) [26]. As expected, Neolithic samples present an excess of genetic contribution to southern Europeans, in particular to modern Sardinians, when compared to Bronze Age samples, which in turn consistently share more haplotypes with northern/eastern groups.

**Fig 3.**
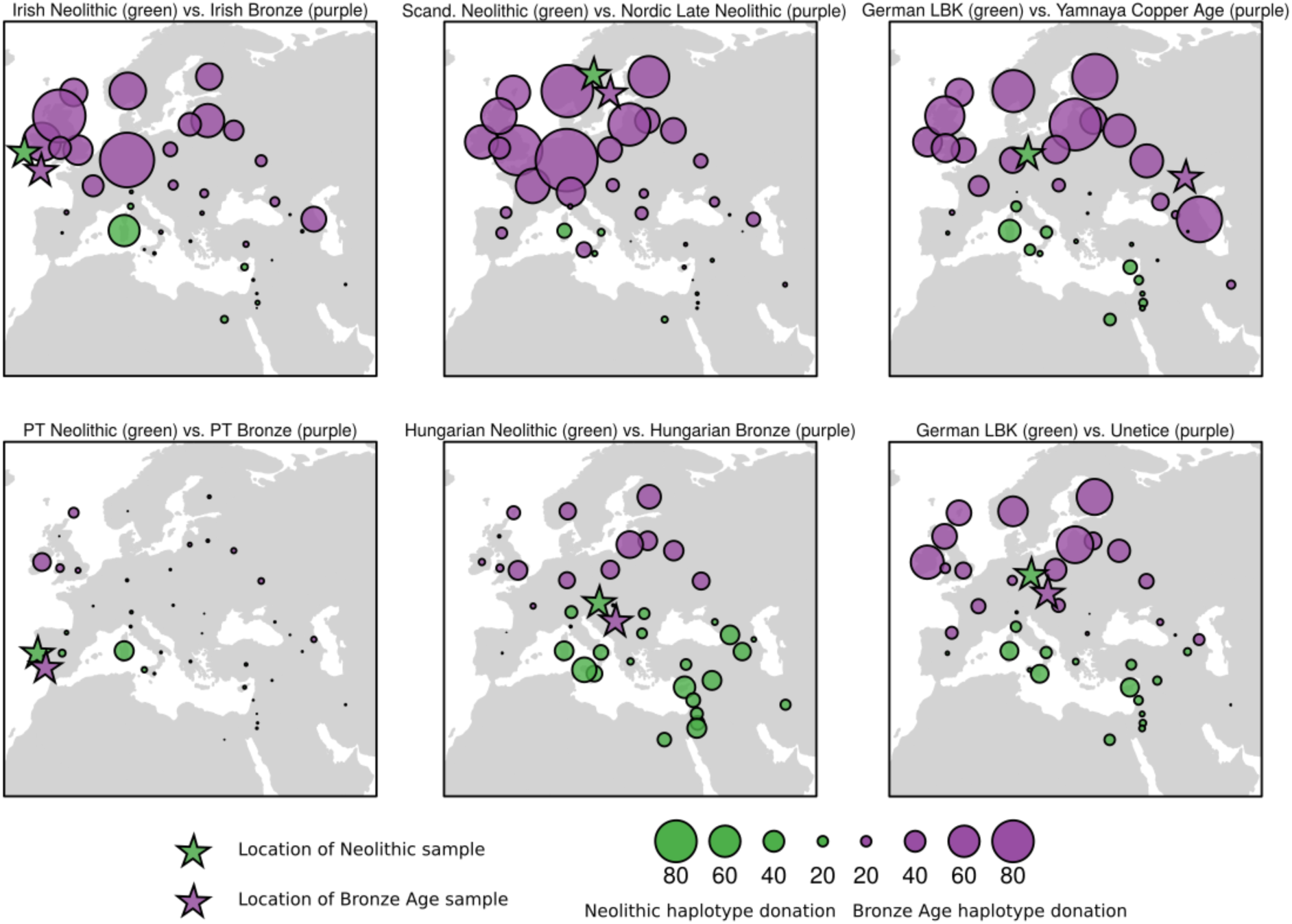
Total Variation Distance between vectors of median haplotype donation from Bronze Age (purple) and Neolithic (green) samples from different regions in Europe to modern populations. Circle size varies according to the absolute difference between Neolithic and Bronze Age samples in terms of the number of haplotypes donated to present day populations. Regardless of the geographical locations of the ancient samples, Neolithic samples tend to donate comparatively more haplotypes to Southern populations, while Bronze Age show the opposite pattern, with an excess of haplotype contribution to Northern Europeans. This pattern is present, but distinctly weaker in the Portuguese Neolithic-Bronze Age comparison.

Consistent with this, when comparing Portuguese Neolithic to Bronze Age samples, the former presented an excess of haplotype donation to Sardinian and Spanish (p=0.017). Northern/eastern ancestry is evident in the Bronze Age, with significantly increased enrichment in Chuvash, Orcadian (p=0.017), Lezgin and Irish (p=0.033). However, this shift from southern to northern affinity is markedly weaker than that observed between Neolithic and Bronze Age genomes in Ireland, Scandinavia, Hungary and Central Europe. These findings suggest detectable, but comparatively modest, Steppe-related introgression present at the Portuguese Bronze Age.

### Comparison of ancient samples from Portugal with ancient and modern individuals using directly observed haploid genotypes

#### Bronze Age Y-Chromosome discontinuity

Previous studies have demonstrated a substantial turnover in Y-chromosome lineages during the Northern European Late Neolithic and Bronze Age, with R1b haplogroup sweeping to high frequencies. This has been linked to third millennium population migrations into Northern Europe from the Steppe, hypothesised to have introduced Indo-european languages to the continent [4]. Strikingly, the array of Y-chromosome haplotypes in ancient Iberia shifts from those typical of Neolithic populations to haplogroup R1b-M269 in each of the three BA males. Interestingly, modern Basque populations have this variant at high frequency (87.1%) [12].

#### ADMIXTURE analysis and D-Statistics

ADMIXTURE analysis of the Portuguese with a wider array of modern and ancient samples was possible using pseudo-diploid calls and allowed us to visualise the temporal and geographical distribution of the major European ancestral components (Fig 4). An increase in the dominant ancestral coefficient of European HG individuals (coloured red) is clear between early and subsequent Iberian Neolithic populations but no discernable difference in HG ancestry is visible between Portuguese MN individuals on the Atlantic coast and their contemporaries from Northeast Spain, suggesting similar admixture processes [4,17]. This increase in WHG admixture in Portuguese MN and LNCA relative to an earlier Cardial Neolithic is also detectable through *D*-Statistic tests (S4 Text; S6 Table), with WHG from Spain, Switzerland and Luxembourg yielding higher levels of significance in comparison the Hungarian WHG KO1 for the test D(Mbuti, WHG; Cardial, MN/LNCA), supporting fineSTRUCTURE results. *D*-Statistics also revealed both the Portuguese MN and LNCA individuals to share higher affinity to Early Neolithic samples from Spain and Greece over Hungarian, LBK and NW Anatolian groups. The Portuguese MN and LN formed clades with each other to the exclusion of all other groups tested, suggesting some level of regional continuity across the Middle to Late Neolithic of Portugal.

**Fig 4.**
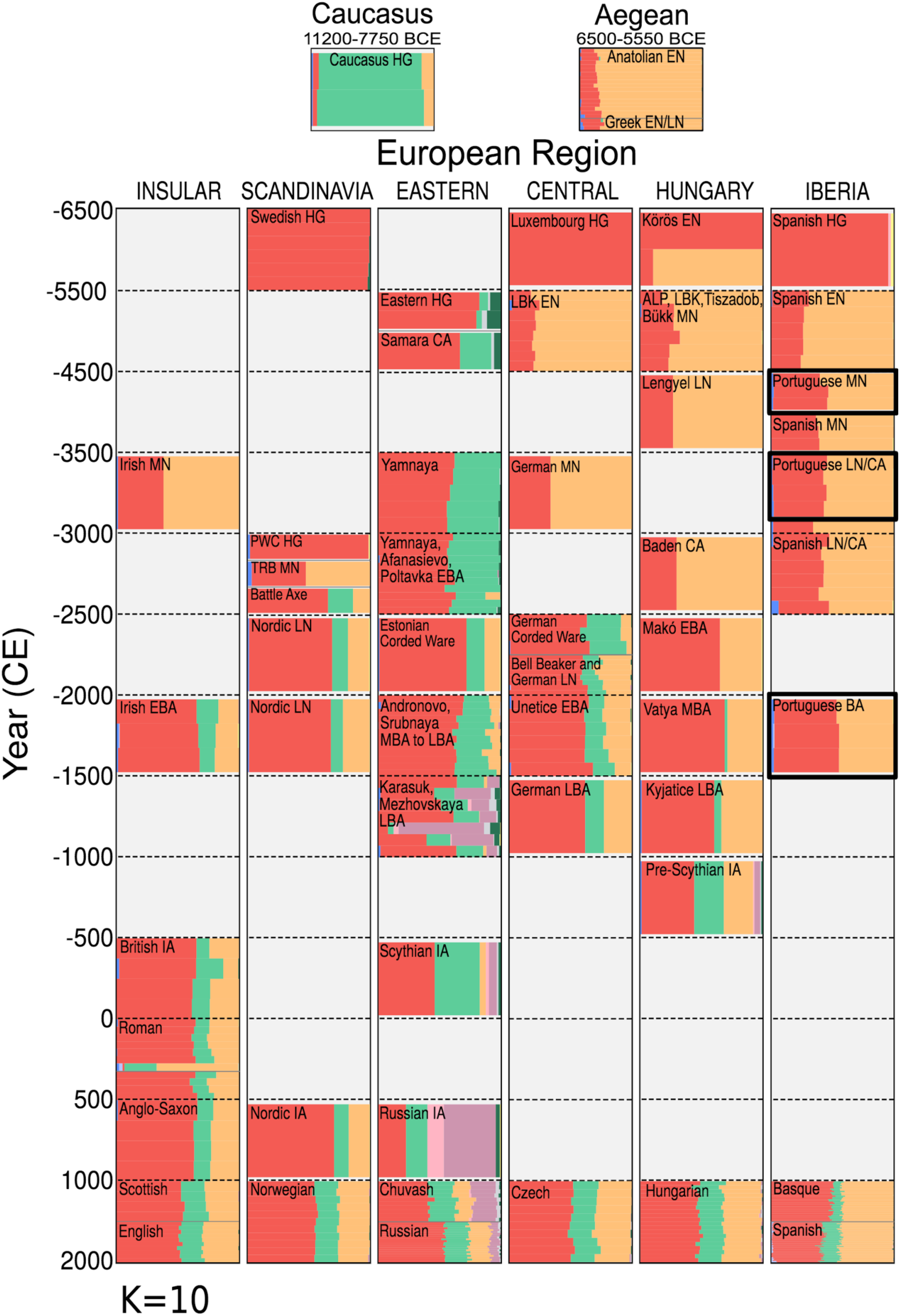
ADMIXTURE analysis of 1941 modern and 176 ancient individuals. Selected profiles of 169 ancient samples, alongside individuals from nine present-day Eurasian populations are displayed here for K=10 ancestral clusters. Individuals are ordered within a grid, partitioned by approximate time period and geographic region. Where possible, ancient individuals have been grouped under common population labels, based on archaeological context. For populations containing three or less individuals, bar plots have been narrowed, leaving empty space within the grid box. Samples from the current study are highlighted in bold.

A recurring feature of ADMIXTURE analyses of ancient northern Europeans is the appearance and subsequent dissemination within the Bronze Age of a component (teal) that is earliest identified in our dataset in HGs from the Caucasus (CHG). Unlike contemporaries elsewhere (but similarly to earlier Hungarian BA), Portuguese BA individuals show no signal of this component, although a slight but discernible increase in European HG ancestry (red component) is apparent. *D*-Statistic tests would suggest this increase is associated not with Western HG ancestry, but instead reveal significant introgression from several steppe populations into the Portuguese BA relative to the preceding LNCA (S4 Text, S6 Table). Interestingly, the CHG component in ADMIXTURE is present in modern-day Spaniards and to a lesser extent in the Basque population.

### Polygenic risk score analysis of height in ancient samples

Height can be expected to give the most reliable predictions due its strong heritability and massive scale of genome wide association studies; the GIANT consortium has estimated 60% of genetic variation as described by common variants [27]. Using the imputed data of >500 thousand diploid SNP calls [27] we combined genetic effects across the whole genome to estimate this phenotype in individuals. Fig 5 plots genetic height in ancient individuals and reveals clear temporal trends. European hunter-gatherers were genetically tall and a dramatic decrease in genetic height is associated with the transition to agriculture (p<0.001). During the Neolithic period we see a steady increase, probably influenced by admixture with hunter-gatherers. Within this trend, Iberian individuals are typical of the Middle and Late Neolithic and we see no evidence of an Iberian-specific diminution as has been previously suggested from a 180 SNP panel [24] (Fig 5; S7 Text, S26 Fig). This increase continues through the Bronze Age, influenced in part by admixture with Steppe introgressors who have high predicted values (Neolithic vs *Yamnaya_Afanasievo*, p<0.018) and into the early centuries AD where ancient Britons and Anglo-Saxons are among the tallest in the sample. That Yamnaya and hunter-gatherer introgressions are major determinant of height variation is supported by strong correlations between these ancestral components and genetic height in modern European populations (Fig 4, S7 Text, S27 Fig).

**Fig 5.**
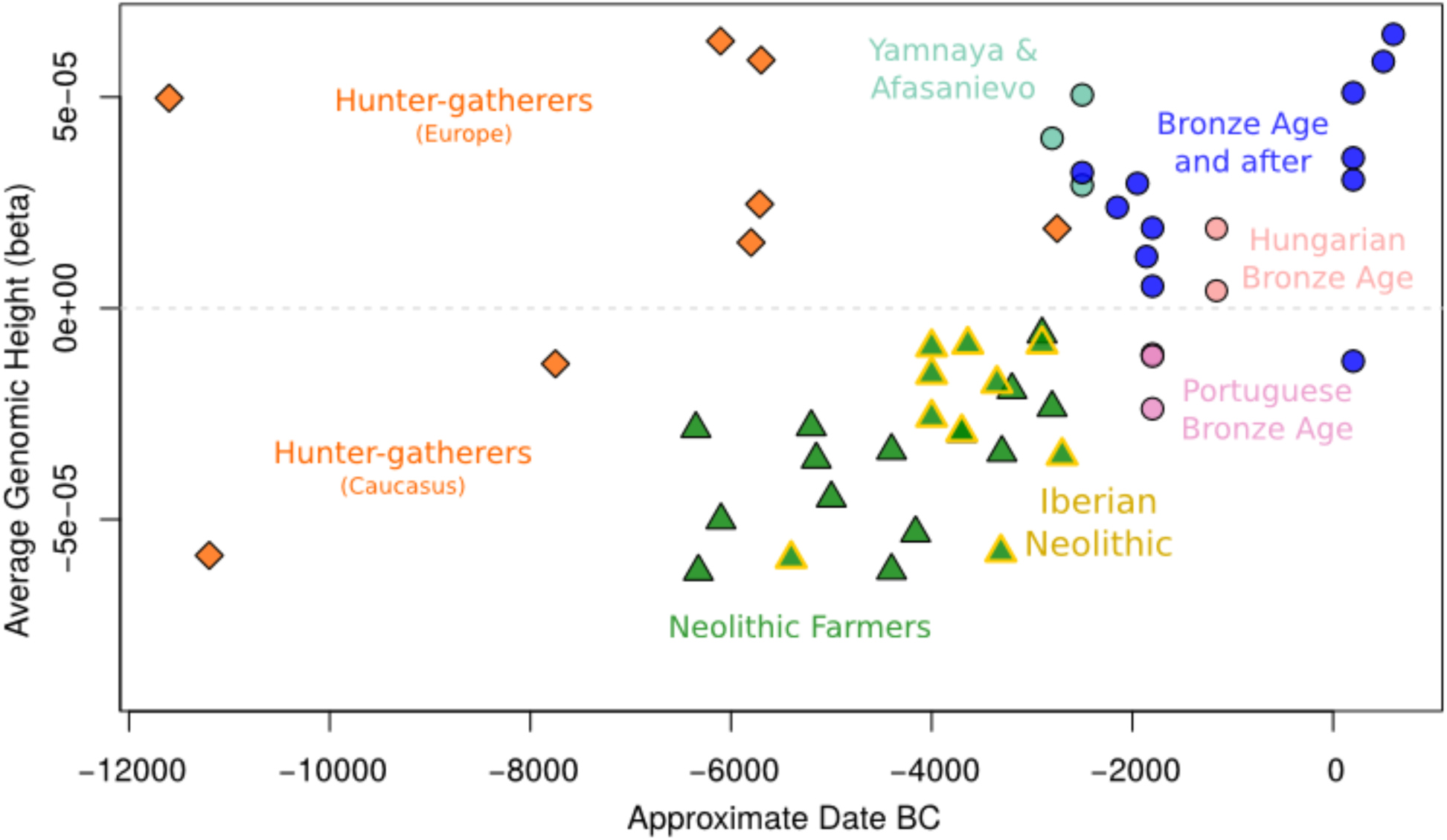
Average genomic height for each of the Western Eurasian samples in the imputed dataset, plotted against its approximate date, highlighting temporal trends in genetic height. We excluded from this analysis Russian Bronze and Iron Age individuals containing variable amounts of Siberian admixture, but polygenic scores for all imputed samples can be seen in S7 Text.

### Extended Haplotype Homozygosity

The role of positive selection in shaping diversity at specific loci in European populations has been of enduring interest and thus we tested whether our imputed genomes could directly reveal the imprint of adaptation in the past. For this we used the extended haplotype homozygosity method [28] with the six loci related to diet and pigmentation highlighted in the analysis by [24]: LCT (rs4988235), SLC24A5 (rs1426654), SLC45A2 (rs16891982), HERC2 (rs12913832), EDAR (rs3827760) and FADS1 (rs174546) (S8 Text, S32 Fig). Two of these, LCT and FADS1 showed strong signals consistent with selective sweeps; homozygous haplotypes that are longer than those surrounding the derived selected allele and that are also markedly longer than those observed in modern populations (Fig 6). The selective sweep signal for LCT (driven by adaptation to a dietary reliance on raw milk) appears in the Bronze Age and that associated with FADS1 shows first in the Neolithic sample, supporting that this may be a response to changes in the spectrum of fatty acid intake afforded after the transition to an agricultural diet [29].

**Fig 6.**
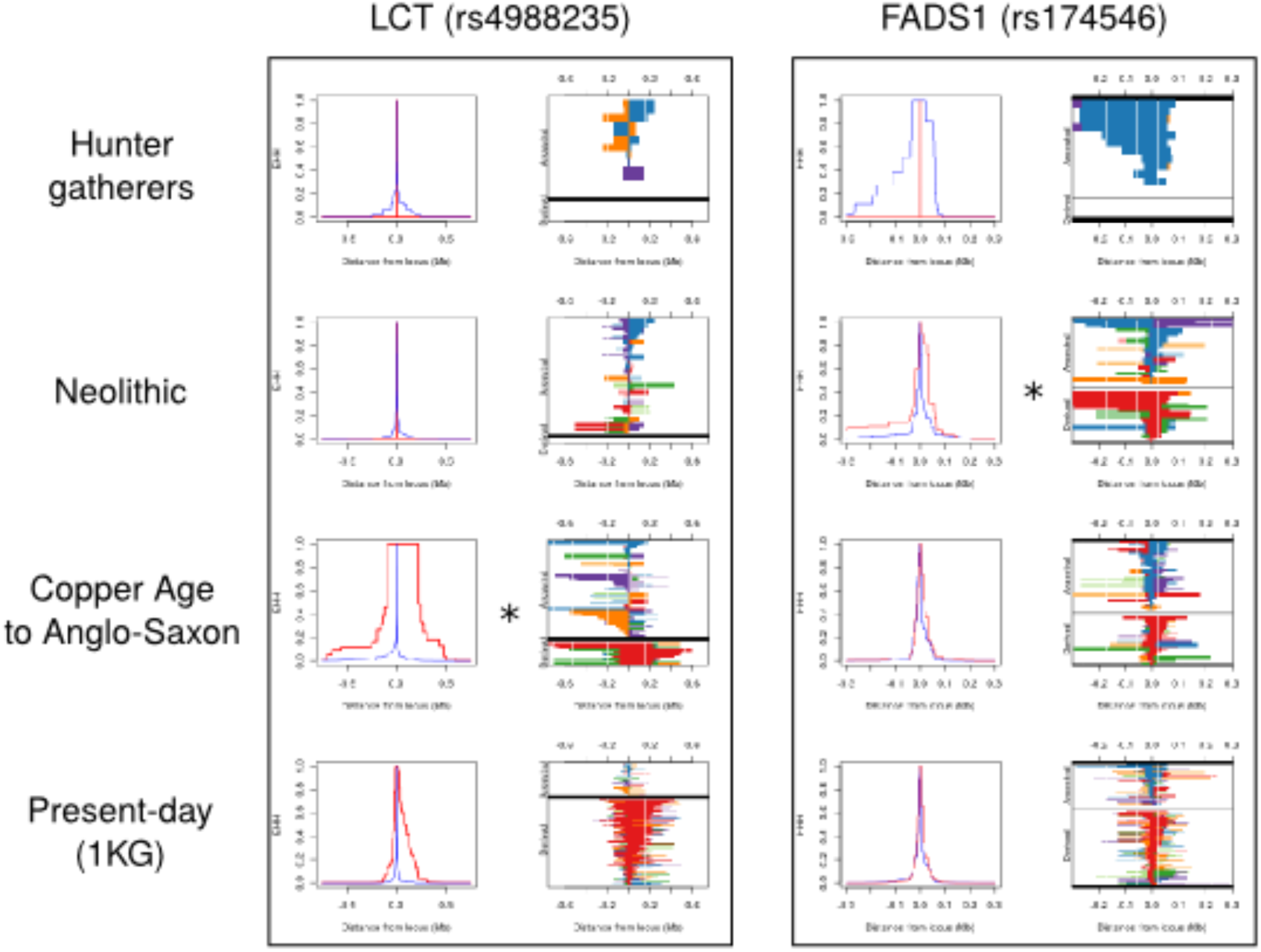
Extended haplotype homozygosity in regions under selection. Panels on the left represent the decay of EHH, or the probability of homozygosity at a certain base across 2 randomly chosen chromosomes in a population. Plots on the right represent existing haplotypes in a population, with the lower portion of the graph depicting haplotypes with the derived allele (red) and the upper part showing haplotypes carrying the ancestral allele (blue). Unique haplotypes in a population are not represented. Legend: CEU -Utah Residents (CEPH) with Northern and Western Ancestry; YRI -Yoruba in Ibadan, Nigeria; CHB -Han Chinese in Beijing, China; 1KG: 1000 Genomes Project. * Earliest appearance of the homozygous derived allele in the samples analysed.

## Discussion

Our genomic data from 14 ancient individuals from 8 Portuguese archaeological contexts ranging from the Middle Neolithic to Middle Bronze Age throws light on how the two fundamental transitions in European prehistory affected populations at the Atlantic edge. Previous data from north Mediterranean regions in Iberia have shown that the first farmers had predominantly Anatolian ancestry [4,18,21], with some increase in hunter-gatherer admixture occurring between the Early and Middle Neolithic. Our analyses, using both observed haploid SNPs and imputed diploid haplotypes show this pattern extends to the Atlantic coast of the peninsula, a region where a dense Mesolithic population persisted in the Neolithic for some 500 years. We support Middle Neolithic HG admixture having occurred locally, as there is greater haplotypic affinity of these Iberians to HG genomes from western Europe than to a hunter-gatherer genome excavated from a much earlier point of contact within the spread of the Neolithic; that within a Hungarian settlement representative of the earliest agricultural cultures of southeast Europe. This affinity is not shared by the earlier genome from the classical Neolithic Cardial phase (7500-7100 BP) which supports the geographical adjacency of this Middle Neolithic HG admixture.

Imputation of ancient European genomes sequenced to 1x coverage has been shown to give diploid genotypes at ∼99% accuracy [3]. Our imputation of 67 genomes yielded genome-scale diploid calls which we surmised should allow the prediction of polygenic traits at the individual level. We illustrate this for height, in which combined genomewide locus effects are known to explain a high proportion of trait variance and which has been shown to have been under selection in Europeans [24,30,31]. Most strikingly, we find that European hunter-gatherers are significantly taller than their early Neolithic farming counterparts. A pattern of increasing genetic height with time since the Neolithic is clear in these European individuals, which may be influenced by increasing admixture with populations containing higher ancestral components of Eurasian hunter-gatherers. This concords with the increased forager-farmer admixture in the transition from the Early to Middle Neolithic; including within Iberian Neolithic individuals. Interestingly, this is in contrast to previous results which estimated a height decrease within this group. However, that work used more limited data, 169 predictive loci, and predicted at a population rather than individual level using a minimum of only two chromosomes called per SNP [24]. Genetic height increases through the Bronze Age are further influenced by Yamnaya introgression and continue through to a series of early Britons sampled from the early centuries AD. Within this time frame, the genetically tallest individual is an Anglo-Saxon from Yorkshire, followed by a Nordic Iron Age sample.

Our analyses yield both signals of continuity and change between Portuguese Neolithic and Bronze Age samples. ADMIXTURE analysis showing similar ancestral components, and higher order branching in fineSTRUCTURE clustering suggest a level of continuity within the region. Also, both show a degree of local European HG admixture (relative to central European HG influence) that is not observed within other samples in the data set. However, final fineSTRUCTURE clustering and the PCA plot places the Portuguese BA as a separate group which is intermediate between Atlantic Neolithic samples and the Central European Bronze Age individuals. *D*-statistics support some influx of ancestral elements derived from the east, as is seen in the northern Bronze Age, and a distinct change in Y-chromosome haplotypes is clear -all three Iberian BA males are R1b, the haplogroup that has been strongly associated with Steppe-related migrations. Patterns of haplotype affinity with modern populations illustrate the Portuguese population underwent a shift from southern toward northern affinity to a distinctly reduced degree to that seen with other regional Neolithic-BA transitions.

Taken together this is suggestive of small-scale migration into the Iberian Peninsula which stands in contrast to what has been observed in Northern, Central [4,5] and Northwestern Europe [11] where mass migration of steppe pastoralists during the Copper Age has been implied. The Y-chromosome haplotype turnover, albeit within a small sample, concords with this having been male-mediated introgression, as suggested elsewhere for the BA transition [32].

Several candidate windows for the entry of Steppe ancestry into Portugal exist. The first is the possible emergence of Bell Beaker culture in Southwest Iberia and subsequent establishment of extensive networks with Central and NW European settlements, opening up the possibility of back-migration into Iberia. Indeed, Central European Bell Beaker samples have been observed to possess both steppe-related ancestry and R1b-P312 Y-chromosomes [4,5]. Furthermore, through the analysis of modern samples, it has been proposed that the spread of Western R1b-lineages fits with the temporal range of the Corded Ware and Bell Beaker complexes [33].

An alternative is in the Iberian Middle to Late Bronze Age when individualized burials became widespread and bronze production began [34]. At this time the spread of horse domestication enabled unprecedented mobility and connectedness. This was coupled with the emergence of elites and eventually led to the complete replacement of collective Megalithic burials with single-grave burials and funerary ornamentation reflecting the status of the individual in society. These changes are seen in the Iberian Bronze Age, with the appearance of cist burials and bronze daggers [35]. Indeed, two of the Bronze Age samples analysed in the present work belong to an archaeological site in SW Iberia where the earliest presence of bronze in the region was demonstrated, as well as high status burials with elaborate bronze daggers [35,36].

Two alternate theories for the origin and spread of the Indo-European language family have dominated discourse for over two decades: first that migrating early farmers disseminated a tongue of Neolithic Anatolian origin and second, that the third Millennium migrations from the Steppe imposed a new language throughout Europe [37,38] [4]. Iberia is unusual in harbouring a surviving pre-Indo-European language, Euskera, and inscription evidence at the dawn of history suggests that pre-Indo-European speech prevailed over a majority of its eastern territory with Celtic-related language emerging in the west [39]. Our results showing that predominantly Anatolian-derived ancestry in the Neolithic extended to the Atlantic edge strengthen the suggestion that Euskara is unlikely to be a Mesolithic remnant [17,18]. Also our observed definite, but limited, Bronze Age influx resonates with the incomplete Indo-European linguistic conversion on the peninsula, although there are subsequent genetic changes in Iberia and defining a horizon for language shift is not yet possible. This contrasts with northern Europe which both lacks evidence for earlier language strata and experienced a more profound Bronze Age migration.

## Material and Methods

### Ancient DNA sampling, extractions and sequencing

All ancient DNA (aDNA) work was done in clean-room facilities exclusively dedicated to this purpose at the Smurfit Institute, Trinity College Dublin, Ireland. We extracted DNA from ∼100 mg of temporal bone samples belonging to 14 samples from 8 archaeological sites in Portugal ranging from the Mid Neolithic to the Mid Bronze Age in Portugal (S1 Text) using a silica-column-based method [40] with modifications [41]. We incorporated DNA fragments into NGS libraries using the library preparation method described in [42] and amplified these with 2-4 different indexing primers per samples and purified (Qiagen MinElute PCR Purification Kit, Qiagen, Hilden, Germany) and quantified (Agilent Bioanalyzer 2100). Samples were sequenced to ∼1.15X (0.05-2.95X) in an Illumina HiSeq 2000 (100 cycle kit, single-end reads mode; Macrogen) (S2 Text).

### Read processing and analysis

We used Cutadapt v. 1.3 [43] to trim NGS read adapters and aligned reads to the human reference genome (UCSC hg19) and mtDNA (rCRS, NC_012920.1) with the Burrows-Wheeler Aligner (BWA) v.0.7.5a-r405 [44], filtering by base quality 15, removing PCR duplicates and reads with mapping quality inferior to 30 using SAMtools v.0.1.19-44428cd [45]. We estimated genomic coverage with Qualimap v2.2 [46] using default parameters (S2 Text).

### Contamination Estimates and Authenticity

In order to assess the level of contamination in ancient samples, we considered the number of mismatches in mtDNA haplotype defining mutations and determined the number of X-chromosome polymorphisms in male samples (S2 Text) [47]. We analysed aligned reads using mapDamage v2.0 [48] to inspect patterns of aDNA misincorporations, which confirm the authenticity of our data.

### Sex determination and uniparental lineage determination

We used the method published in reference [49] to determine the sex of the ancient individuals (S2 Text, S3 Fig). Y-chromosome lineages of ancient male samples were identified using Y-haplo software [50] (https://github.com/23andMe/yhaplo, S4 Table). For mtDNA analysis, reads were separately aligned to the revised Cambridge Reference Sequence (rCRS; NC_012920.1) [51], filtering for base (q ≥ 20) and mapping (q ≥ 30) quality and duplicate reads as above. mtDNA haplogroups were identified using mtDNA-server (http://mtdna-server.uibk.ac.at/start.html, with default parameters.

### Comparison with modern and ancient individuals

Smartpca version 10210 from EIGENSOFT [52,53] was used to perform PCA on a subset of West Eurasian populations (604 individuals) from the Human Origins dataset [2], based on approximately 600,000 SNPs (S3 Text, S4 Fig). The genetic variation of 239 ancient Eurasian genomes [2,4,5,11,14–21,24,54–57] was then projected onto the modern PCA (lsqproject: YES option). A model-based clustering approach implemented by ADMIXTURE v.1.23 [58] was used to estimate ancestry components in 10 of the Portuguese samples, alongside 1941 modern humans from populations worldwide [2] and 166 ancient individuals. Only ancient samples with a minimum of 250,000 genotyped markers were included. The dataset was also filtered for related individuals, and for SNPs with genotyping rate below 97.5%. A filter for variants in linkage disequilibrium was applied using the --indep-pairwise option in PLINK v1.90 with the parameters 200, 25 and 0.4. This resulted in a final 219,982 SNPs for analysis. ADMIXTURE was run for all ancestral population numbers from K=2 to K=15, with cross-validation enabled (--cv flag), and replicated for 40 times. The results for the best of these replicates for each value of K, i.e. those with the highest log likelihood, were extremely close to those presented in [11]. The lowest median CV error was obtained for K=10.

### D-statistics

Formal tests of admixture were implemented using D-statistics [59] and F-statistics [60,61] using the AdmixTools package (version 4.1) (Patterson 2012). These were carried out on WGS ancient data only, using autosomal biallelic transversions from the 1000 Genomes phase 3 release (S4 Text, S5 and S6 Table).

### Genotype Imputation

We selected for genotype imputation all published samples that had been sequenced by whole-genome shotgun sequencing and for which coverage is above 0.75X, including 4 ancient individuals downsampled to 2X which were included for estimating accuracy. Within these were called ∼77.8 million variants present in the 1000 Genomes dataset using Genome Analysis Toolkit (GATK) [62], removing potential deamination calls. These were used as input by BEAGLE 4.0 [63] which phased and imputed the data (S5 Text, S7 Table). This resulted in a VCF file with approximately 30 M SNPs.

### fineSTRUCTURE analyses

In analysis I (Fig 1), imputed variants in 67 ancient Eurasian samples were filtered for posterior genotype probability greater or equal to 0.99. Variants not genotyped across all individuals were removed with vcftools [64], also excluding SNPs with MAF < 0.05, resulting in approximately 1.5 M SNPs and the resulting VCF was converted to IMPUTE2 format with bcftools version 0.1.19-96b5f2294a (https://samtools.github.io/bcftools/). Hap files were converted to CHROMOPAINTER format with the script “impute2chromopainter.pl”, available at http://www.paintmychromosomes.com/ and created a recombination map with “makeuniformrecfile.pl”. We then split the dataset by chromosome with vcftools and ran CHROMOPAINTER and fineSTRUCTURE v2 [23] with the following parameters: 3,000,000 burn-in iterations, 1,000,000 MCMC iterations, keeping every 100^th^ sample. In S6 Text we describe all 5 analysis CHROMOPAINTER and fineSTRUCTURE analyses in more detail: I -aDNA samples only (S7-S20 Fig); II -aDNA samples and present-day Eurasians and Yoruba (Fig 3); III -Comparison of linked and unlinked analyses (S21 Fig); IV -Analysis with unfiltered genotype probabilities; V -Detection of biases in CHROMOPAINTER analyses derived from genotype imputation in ancient samples (S22 and S23).

### Polygenic traits in ancient samples

Genetic scores for polygenic traits including height [27], pigmentation [65], Anthropometric BMI [66] and T2D [67] in 67 ancient samples were estimated using PLINK [68] using the --score flag. Odds ratio in the T2D summary statistics [67] were converted to effect size by taking the logarithm of OR/1.81 [69]. In our analyses, we compared p-value filtering (unfiltered p-value threshold against p<0.001) when possible to qualitatively evaluate robustness of signals observed (S7 Text, S25, S30, S31 Fig).

In order to investigate the correlation between ancient ancestry in present-day populations and height genetic scores, we first calculated polygenic risk in Eurasian populations from the Human Origins dataset. This was followed by the estimation of the percentage ancestry of five distinct ancient populations (EHG, CHG, WHG, Yamnaya, Anatolian Neolithic) in the same dataset, which was done through the implementation of the F4 ratio method described in [61] using the Admixtools package (version 4.1). Two individuals possessing the highest genomic coverage from each population were used in the test, which took the form f4(Mbuti, Ancient_Ind1; Modern_WEurasinan, Dai)/f4(Mbuti, Ancient_Ind1; Ancient_Ind2, Dai) (S8 Table).

### Extended Haplotype Homozygosity analysis

We used Selscan [70] to investigate extended haplotype homozygosity (EHH) around SNPs of interest previously described in [71]: LCT (rs4988235), SLC24A5 (rs1426654), SLC45A2 (rs16891982), HERC2 (rs12913832), EDAR (rs3827760) and FADS1 (rs174546). First, SNPs within 5 Mb of each SNP were included for analysis, removing SNPs which are multiallelic and with multiple physical coordinates. EHH requires large populations, and therefore we used selscan in 3 groups: HG, Neolithic farmers and Copper Age to Anglo-Saxon, using the --ehh and --keep-low-freq flag (S8 Text).

## Acknowledgements

The authors wish to acknowledge the DJEI/DES/SFI/HEA Irish Centre for High-End Computing (ICHEC) for the provision of computational facilities and support.

## Web resources

Raw data and aligned reads have been submitted to http://www.ebi.ac.uk/ena/data/view/PRJEB14737, secondary accession ERP016408.

## Competing interests

The authors declare no conflict of interest.

## Supporting Information captions

S1 Text. Archaeological information.

S2 Text. Ancient DNA analysis.

S3 Text. Comparison of ancient samples with other ancient and modern datasets using genotype data.

S4 Text. Exploring ancient Iberian affinities through F-and D-statistics.

S5 Text. Imputation of missing genotypes in ancient samples.

S6 Text. CHROMOPAINTER and fineSTRUCTURE analyses.

S7 Text. Analysis of polygenic traits.

S8 Text. Extended Haplotype Homozygosity Analysis.

S1 Table -X-chromosome contamination estimated with ANGSD (Korneliussen et al. 2014) and based on a previously published method (Rasmussen et al. 2011).

S2 Table - X-chromosome contamination based on the number of mismatches at X-chromosome SNPs and adjacent sites.

S3 Table - mtDNA lineages and contamination estimates based on mismatches at haplotype defining sites.

S4 Table - Y-chromosome lineages determined in the ancient Portuguese samples.

S5 Table - D-statistics in the form of D(Mbuti, X; Y, Z) to test admixture between ancient populations.

S6 Table - Selected D-statistics associated with Portuguese Neolithic and Bronze samples.

S7 Table - List of ancient samples selected for genotype imputation.

S8 Table - List of ancient individuals used in the F4 ratio test.

This table contains the individuals which were used to estimate the approximate percentage ancestry in modern populations of five ancestral groups who have contributed to western Eurasian variation, using an F4 ratio test (Patterson 2012).

S1 Fig - Map and geographical locations of the archaeological locations of the samples sequenced in the present study.

S2 Fig - Sex Determination using Ry_compute.

S3 Fig - Post-mortem misincorporations in ancient samples.

S4 Fig - Principal component analysis of 604 modern West Eurasians onto which variation from 224 ancient genomes has been projected.

The analysis is based on approximately 600,000 SNP positions. Moderns samples from the Human Origins dataset are represented in greyscale, with the exception of modern Iberians shown in green. Ancient samples are coloured by time depth and shaped according to geographic region. Ancient individuals from Portugal are outlined in red.

S5 Fig -Estimation of Imputation accuracy on chromosome 21.

Comparison of variant calls obtained for BR2, NE1, Loschbour and Stuttgart at full coverage with genotypes from the same 4 individuals downsampled to 2.5x and subsequently imputed. Accuracy in (A) all 3 types of genotypes; (B) homozygous reference; (C) heterozygous and (D) homozygous alternate.

S6 Fig - Proportion of correctly imputed genotypes grouped by minor allele frequency bins of 0.005.

In this analysis, imputed genotypes were filtered by post imputation genotype probability ⋝ 0.99.

S7 Fig - Geographical and PC genetic coordinates for the Western_HG1 cluster.

S8 Fig - Geographical and PC genetic coordinates for the Western_HG2 cluster.

S9 Fig - Geographical and PC genetic coordinates for the fineSTRUCTURE Scandinavian_HG cluster.

S10 Fig - Geographical and PC genetic coordinates for the fineSTRUCTURE cluster Caucasus Hunter-gatherers.

S11 Fig - Geographical and PC genetic coordinates for the fineSTRUCTURE AegeanEN_HungarianLBK cluster.

S12 Fig - Geographical and PC genetic coordinates for the fineSTRUCTURE HungarianMLN_SpainCardialEN cluster.

S13 Fig - Geographical and PC genetic coordinates for the fineSTRUCTURE Atlantic_Neolithic cluster.

S14 Fig - Geographical and PC genetic coordinates for the fineSTRUCTURE Yamnaya_Afanasievo cluster.

S15 Fig - Geographical and PC genetic coordinates for the fineSTRUCTURE Sintashta_Andronovo cluster.

S16 Fig - Geographical and PC genetic coordinates for the fineSTRUCTURE CopperAge_to_AngloSaxon cluster.

S17 Fig - Geographical and PC genetic coordinates for the fineSTRUCTURE Hungary_BA cluster.

S18 Fig - Geographical and PC genetic coordinates for the fineSTRUCTURE Portugal_BA cluster.

S19 Fig - Geographical and PC genetic coordinates for the fineSTRUCTURE Russia_LBA cluster.

S20 Fig - Geographical and PC genetic coordinates for the fineSTRUCTURE Russia_LBA_IA cluster.

S21 Fig - Comparison between (A) unlinked and (B) linked CHROMOPAINTER/fineSTRUCTURE analyses.

The unlinked analysis is only able to identify 10 populations, 9 less than when incorporating the linkage model.

S22 Fig -CHROMOPAINTER haplotype donation vectors between each one of the imputed and non-imputed samples

(A) Correlation between imputed and non-imputed median haplotype donation from sample BR2 (1), Loschbour (2) and LBK (3). (B) Normal Quantile-Quantile plots and outlier detection (labelled populations). Coloured dots show populations present (red) or absent (black) in the 1000 Genomes reference haplotype dataset. (C) Barplots illustrating imputed (left) and non-imputed (right) median haplotype donation (light blue) and the difference between median haplotype donation per population (dark blue).

S23 Fig - fineSTRUCTURE tree comparison between each one of the imputed and non-imputed samples (BR2, Loschbour and LBK).

The position of aDNA samples (shown in red) is very similar in both analyses.

S24 Fig - Bar plots illustrating polygenic risk scores across time, estimated for each one of the ancient population clusters.

The traits chosen were: A) Height; B) Pigmentation; C) BMI and D) T2D. Polygenic scores were centered at the mean for the dataset. As in Fig 1 in the main text, each cluster is represented with a different colour.

S25 Fig -Polygenic risk scores estimated for height using genomewide summary statistics from the Wood 2014 dataset.

(A) p=0 (B) p<0.001. SNPs with posterior genotype probability of less than 0.99 were excluded from analysis.

S26 Fig - Polygenic risk scores estimated for height using genomewide summary statistics.

S27 Fig - Correlation between strands of ancestry and inferred polygenic risk score in present-day Europeans.

Hunter-gatherer (WHG, EHG, CHG), Neolithic (Anatolian_EN) and Steppe (Yamnaya) Ancestry was measured by f4(Mbuti, Ancient_Ind1; Modern_WEurasian, Dai)/f4(Mbuti, Ancient_Ind1; Ancient_Ind2, Dai). Polygenic risk scores for height (92) were determined using ∼280.000 SNPs in 48 European populations. Blue line presents the linear regression. Individual samples are represented by gray dots and larger coloured circles represent the mean genetic score for each population.

S28 Fig - Height map and PCA.

Red - increased genetic height scores, black - decreased genetic height. Broadly, hunter-gatherers and populations from Copper age and after present highest proportion of height increasing associated variants followed by Neolithic farmers.

S29 Fig - Polygenic scores for pigmentation based on the SNPs reported in (94).

SNPs with posterior genotype probability of less than 0.99 were excluded from analysis.

S30 Fig - Polygenic risk scores estimated for BMI using genomewide summary statistics reported by (95) on a multi-ancestry cohort.

(A) p=0 (B) p<0.001. SNPs with posterior genotype probability of less than 0.99 were excluded from analysis.

S31 Fig - Polygenic risk scores estimated for T2D using genomewide summary statistics reported by (96).

A) p=0 B) p<0.001. SNPs with posterior genotype probability of less than 0.99 were excluded from analysis.

S32 Fig - Extended haplotype homozygosity (EHH) in regions under selection. Panels on the left represent the decay of EHH, or the probability of homozygosity at a certain base across 2 randomly chosen chromosomes in a population. Plots on the right represent existing haplotypes in a population, with the lower portion of the graph depicting haplotypes with the derived allele (red) and the upper part showing haplotypes carrying the ancestral allele (blue). Unique haplotypes in a population are not represented. Legend: CEU - Utah Residents (CEPH) with Northern and Western Ancestry; YRI - Yoruba in Ibadan, Nigeria; CHB -Han Chinese in Beijing, China; 1KG: 1000 Genomes Project. * Earliest appearance of the homozygous derived allele in the samples analysed.

